# Changes in the diversity and composition of tree-related microhabitats across climate and human impact gradients on a tropical mountain

**DOI:** 10.1101/2024.05.28.595532

**Authors:** Giovanni Bianco, Andreas Hemp, Matthias Schleuning

## Abstract

Tree-related microhabitats (TReMs) have been proposed as important indicators of biodiversity to guide forest management. However, their application has been limited mostly to temperate ecosystems, and it is largely unknown how the diversity of TReMs varies along environmental gradients. In this study, we assessed the diversity of TReMs on 180 individual trees and 44 plots alongside a large environmental gradient on Kilimanjaro, Tanzania. We used a typology adjusted to tropical ecosystems and a tree-climbing protocol to obtain quantitative information on TreMs on large trees and dense canopies. We computed the diversity of TReMs for each individual tree and plot and tested how TReM diversity was associated with properties of individual trees and environmental conditions in terms of climate and human impact. We further used non-metric multidimensional scaling (NMDS) to investigate the composition of TReM assemblages alongside the environmental gradients. We found that diameter at breast height (DBH) and height of the first branch were the most important determinants of TReM diversity on individual trees, with higher DBH and lower first branch height promoting TReM diversity. At the plot level, we found that TReM diversity increased with mean annual temperature and decreased with human impact. The composition of TReMs showed high turnover across ecosystem types, with a stark difference between forest and non-forest ecosystems. Climate and the intensity of human impact were associated with TReM composition. Our study is a first test of how TReM diversity and composition vary along environmental gradients in tropical ecosystems. The importance of tree size and architecture in fostering microhabitat diversity underlines the importance of large veteran trees in tropical ecosystems. Because diversity and composition of TReMs are sensitive to climate and land-use effects, our study suggests that TReMs can be used to efficiently monitor consequences of global change for tropical biodiversity.

## 1. Introduction

Forest ecosystems are home to two-thirds of the world’s terrestrial biodiversity (Pillay *et al*. 2022) and are among the most threatened ecosystems from anthropogenic land-use change, with the majority of forest ecosystems being degraded to some extent (Watson *et al*. 2018). This makes monitoring forest biodiversity a priority amidst the global biodiversity crisis. However, monitoring biodiversity in forests is time consuming and often difficult, especially for understudied taxa (Hochkirch *et al*. 2021) and in highly diverse ecosystems (Schmeller *et al*. 2017). To address this shortfall, several cost-effective indicators have been devised to indirectly estimate forest biodiversity. Most of these indicators, such as the vertical stratification of vegetation or deadwood availability, aim to describe the capacity of a forest to provide different microhabitats to species (Gao *et al*. 2014). In line with this, the habitat heterogeneity hypothesis states that heterogeneity in biotic and abiotic environmental conditions promotes a higher niche dimensionality, fosters species coexistence, and increases biodiversity (Stein *et al*. 2014). Such patterns appear to hold at different spatial scales (Stein et al., 2014) and have been observed for various taxa (Freemark & Merriam 1986; Price *et al*. 2010), although different organism groups may differ in how they respond to heterogeneity (Heidrich *et al*. 2020).

One of the more recently devised indicator systems of forest biodiversity are tree-related microhabitats (TReMs). TReMs are defined as all structures occurring on a living or dead standing tree, which provide niches to specific organisms at some point of their life cycle. They include structural features of trees like cavities (Fig. 2), bark pockets and damaged branches, but also epiphytic organisms like ferns, mosses, and lichens (Larrieu *et al*. 2018). Given their relevance for many organism groups, TReMs provide a suitable indicator system to quantify the potential occurrence of multiple species simultaneously, they are relatively cost-effective and can be monitored all year round (Asbeck *et al*., 2021). First evidence of TReMs being a reliable predictor of biodiversity comes from studies on bat and insect diversity (Basile *et al*. 2020) and bird diversity (Paillet *et al*. 2018), but these studies have been restricted to temperate forests.

Understanding the determinants of TReM abundance and diversity can help identify important attributes of forest ecosystems and evaluate their role as indicators of environmental change. Studies carried out in temperate forests have concluded that large trees provide more microhabitats (Asbeck et al., 2021). In addition, tree architecture can influence the formation rate of some microhabitats, e.g., by creating weak spots in the crown (Larrieu *et al*. 2022). On a larger scale, climate is known to affect the presence of epiphytes (Benzing 1998) and cavity formation (Remm & Lõhmus 2011) and managed forests generally host fewer TReMs than old-growth forests (Asbeck et al., 2021). So far, most assessments of TReMs have been conducted in temperate and boreal forests, mostly in North America and Europe (Martin *et al*. 2022) and, thus, were restricted to a relatively small pool of tree species (Mamadashvili *et al*. 2023) and a limited range of environmental conditions. This limits our understanding of how TReMs are shaped by climatic factors and human impacts (Martin *et al*. 2022). Studying TReMs along environmental gradients is, thus, essential to understanding how climate and human impact affect TReM diversity and composition.

In this study, we present a first comprehensive assessment of TReMs along a large environmental gradient comprising ten ecosystem types studied along an elevational gradient of more than 3000 meters on Kilimanjaro, Tanzania. We devised a protocol for assessing TReM diversity which allows to record accurate information on TReM presence and abundance even in complex rainforest canopies. Based on this assessment, we analysed the diversity and composition of TReMs on 180 individual trees growing on 44 plots that varied widely in climatic conditions and intensity of human impact. With these data, we tested the following hypotheses:

Larger tree size and more complex tree architecture promote TReM diversity (Larrieu et al. 2022).
TReM diversity at the plot level increases with increasing rainfall and temperature, while it decreases with higher levels of human impact (Asbeck et al., 2021).
Ecosystems characterized by similar environmental conditions are expected to host similar assemblages of TReMs (Asbeck *et al*. 2019).

## 2. Methods

### 2.1 Study Area

Kilimanjaro is a free-standing dormant volcano located in Tanzania (2°45’ – 3° 25’ S, 37° 00’ - 37 43’ E) rising from a plateau at 700m above mean sea level (AMSL) to its summit at 5895m AMSL. Alongside this elevational gradient, the mountain hosts a variety of ecosystems which vary widely in climatic conditions and in intensity of human impacts. Below 1800m AMSL, maize plantations, managed grasslands, coffee plantations and traditional homegardens are the ecosystems with the strongest human footprint, while lowland savannas and lower montane forest are mostly natural ecosystems. Above 1800m, lower montane forest is replaced by *Ocotea* forest, and the latter is then replaced by *Podocarpus* and *Erica* forests at higher elevations. Some *Ocotea* forests have historically experienced selective logging, while *Podocarpus* and *Erica* forests can be damaged by human-caused fires (Hemp 2006).

### 2.2 Study design

We conducted our TReM surveys across three field seasons that took place between February-April 2022, August-November 2022, and January-April 2023. We assessed TReM diversity across 44 plots (50x50 m in size), distributed alongside five elevational transects and representing the ten ecosystem types which harbor trees on the mountain (Figure 1a). These plots were established within the KiLi (FOR 1246) and the Kili-SES (FOR 5064) DFG research units. In these plots, we selected the five largest trees that were deemed safest to climb. Where several options were available, we prioritised sampling trees of different species.

**Figure 1.**
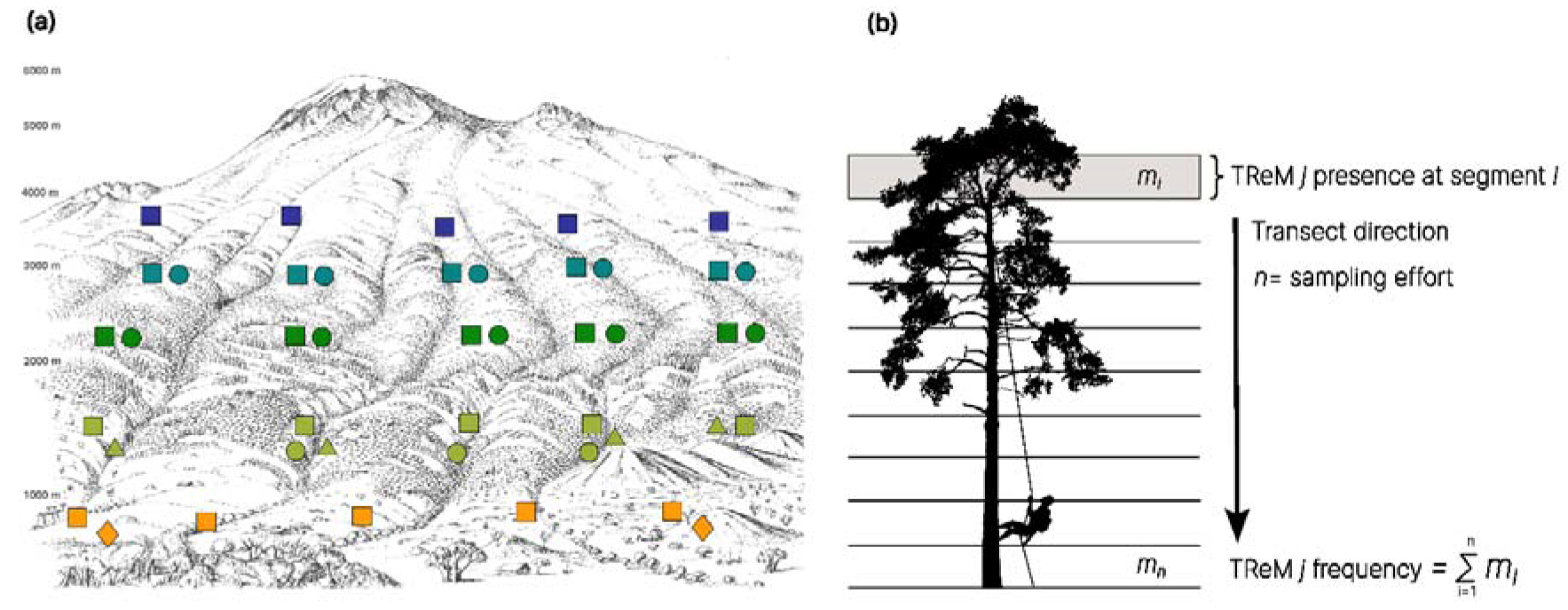
**a) The study system on Kilimanjaro.** The diagram portraits the southern slope of Kilimanjaro and the spatial distribution of the studied plots (n = 44). Orange squares represent natural savanna plots; orange diamonds represent cultivated maize plots. Green circles represent homegardens; green triangles represent coffee plantations; green squares represent lower montane forest. Dark green squares represent *Ocotea* forest and circles disturbed *Ocotea* forest. Aquamarine squares represent *Podocarpus* forest, circles represent disturbed *Podocarpus* forest. Blue squares represent *Erica* forest. Drawing by A. Hemp. **b) Scheme of the tree-climbing survey of TReMs**. The scheme depicts the climber doing the survey descending from the highest anchor point in the tree canopy. The climber records the presence of all 44 TReM types at every one-meter segment. The frequency of each TReM type is given by the sum of the presences for that tree. The sampling effort equals the number of segments surveyed on each tree.

### 2.3 TReM survey

We adapted the hierarchical typology for tree related microhabitats developed by Larrieu et al. (2018) for tropical forests (Nußer *et al*. 2024). To this end, we merged the three kinds of woodpecker breeding cavities into one single TReM type, which encompasses the breeding cavities of woodpeckers, barbets and tinkerbirds (van der Hoek *et al*. 2017). We added epiphytic orchids as an additional TReM type within the epiphyte TReM form because they are known to be fundamental for the life cycle of several pollinator species (Spicer & Woods 2022). We grouped epiphytic vascular plants, which were not orchids, ferns or vines, in the TReM type “other epiphyte”. We further described a new TReM type, named “dead leaves frill”. On Kilimanjaro, this TReM type is specific to plants of the *Dendrosenecio* genus, as these plants build an insulating layer of leaves that protect them from frost and provide shelter to invertebrates (Beck 1986; Tomlinson 1985). A similar structure is also found on plants of the *Espeletia* and *Puya* genera in the Neotropics (Smith 1979). The typology used in this study comprises 44 TReM types corresponding to 6 TReM forms (Cavities, Injuries, Exudates, Deadwood, Fungi and Epiphytes & Epiphytic structures; see Figure 2 for some examples).

**Figure 2.**
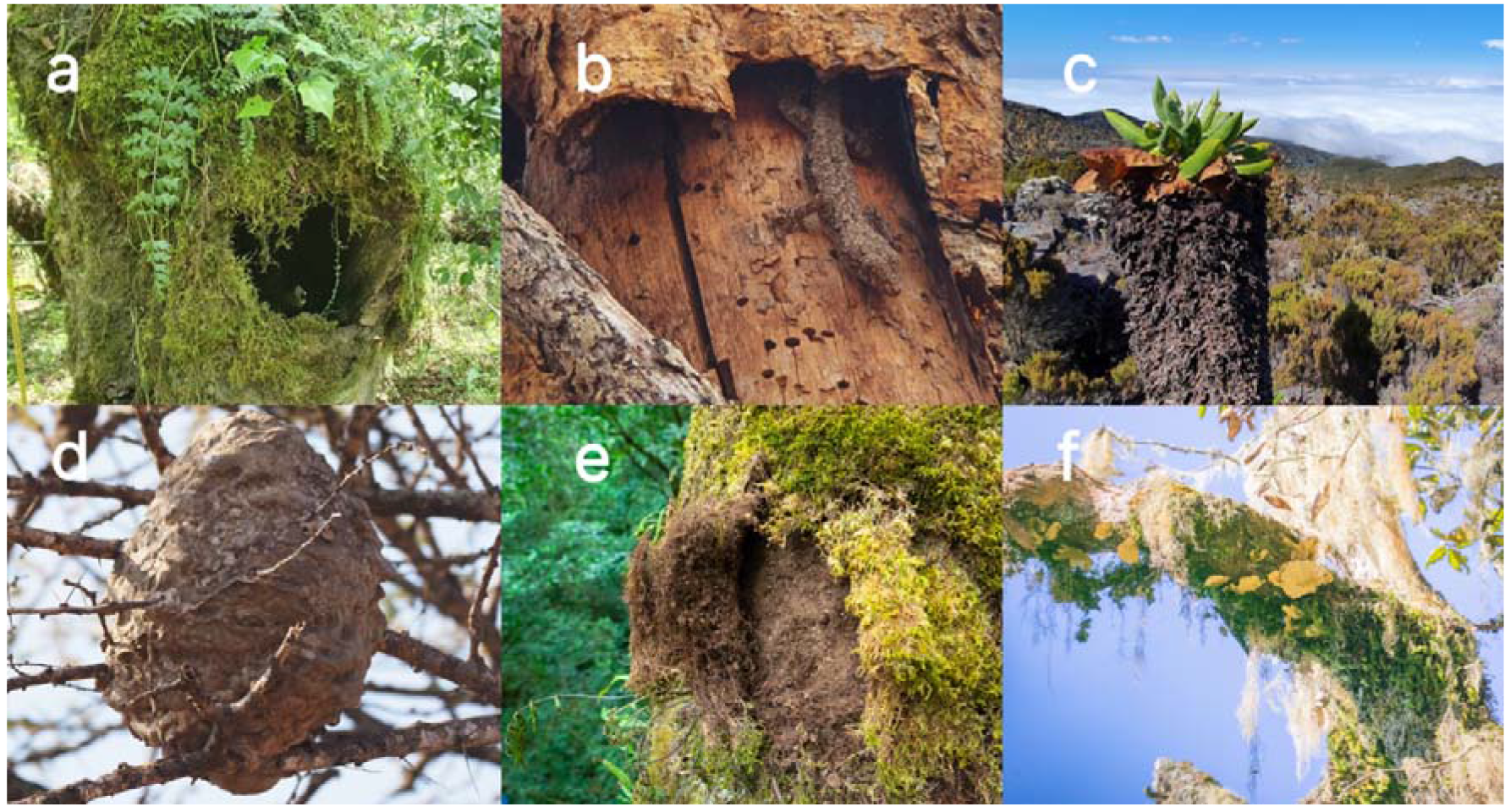
Some of the sampled TReMs on Kilimanjaro. A) Trunk rot-hole. B) Bark shelter, with gecko. C) Dead leaves frill found on *Dendrosenecio spp.* D) Invertebrate nest. E) Bark microsoil forming beneath epiphytic mosses. F) Polypore growing on a dead limb.

For trees taller than five meters, we employed rope-based techniques to access the tree canopy. This was necessary because the tall and complex canopies of tropical forests are difficult to assess from the forest floor. To this end, we set up a throwline on a suitable anchor point from the ground using a catapult (BigShot catapult by Notch Equipment) and throwbags (10oz and 12oz). The throwline was then used to hoist a climbing rope (Courant Rebel, 11mm) for single rope technique up to the anchor point. The tree climber secured himself to the rope via a Petzl ID’s belay/rappel device and proceeded to ascend the tree until the anchor point. If the situation demanded it, once the anchor point was reached, the tree climber set a second line (Tree Runner 12 mm) to continue ascension with Double Rope Technique (Anderson *et al*. 2015). Once at the highest reachable anchor point, the tree climber proceeded to descend the entirety of the tree. During the descent, the climber (G. Bianco) paused at one-meter intervals to assess the presence of each of the 44 TReM types (Fig. 1b). This survey resulted in an incidence matrix of TReM types for each surveyed tree, where rows represent the one-meter segments screened for the 44 TReM types. Based on these data, we were able to calculate the frequency of each TReM type on each sampled tree and plot.

### 2.4 Tree measurements

Within each plot, all trees with a diameter at breast height (DBH) greater than ten cm were identified to the species level, and individually marked with aluminium tags (Ensslin *et al*. 2015). Measurements of the DBH, height, and height of the first branch were recorded for every tagged individual. DBH was measured with a diameter tape (Forestry Suppliers, USA) at 1.3 m for normally shaped trees and 20 cm below or above when branches or irregular shapes impeded measurement at that height. Tree height was measured from the highest ground level around the stem to standardize measurements taken on slopes. For trees which were strongly buttressed or too big to measure by hand, a laser dendrometer (Criterion RD 1000 with TruPulse 200/200, Centennial, USA) was used to measure the tree above the buttresses and at 1.3 m. Height of the highest visible leaf and height of the first branch were measured using an ultra-sonic hypsometer (Vertex IV Hypsometer, Haglöf, Langsele, Sweden) or a laser rangefinder (TruPulse 200/200). We obtained measurement of the wood density for each tagged species by Ensslin et al. (2015), as wood density may be linked to the frequency and type of damage that a tree experiences in its life history (King *et al*. 2006).

### 2.5 Environmental variables

We selected key variables measured at the plot level to describe the variability in climatic conditions and human impacts on Kilimanjaro (Albrecht *et al*. 2021; Peters *et al*. 2016). We selected mean annual temperature (MAT) and mean annual precipitation (MAP) as the main descriptors of the climatic conditions on the mountain. Air temperature was measured in every plot via a sensor placed at a height of about two meters above ground. MAT was calculated from two years of measurements taken at 5 minutes intervals (Peters *et al*. 2016). MAP was interpolated for every plot, using data collected with a network of 70 rain gauges over a time span of 15 years (Hemp 2006).

We obtained metrics of biomass removal (BRI) and vegetation structure (VSI), both associated with the intensity of human impact from Peters et al. (2019). BRI is a measure of direct human impact on a study plot in terms of plant biomass removed by humans via mowing, cattle grazing, fire events, logging, and firewood collection. Estimates of the percentage of plant biomass removed were taken multiple times per plot and cross-checked with information on land use provided by the local landowners (Peters *et al*. 2019). BRI was calculated as the mean of these estimates so that a BRI value of zero indicates a pristine ecosystem without human impact (i.e., no biomass removal), while a value of one indicates that the entirety of plant biomass was removed. VSI quantifies human modification of the vegetation structure relative to the natural conditions at that elevation. To estimate VSI, canopy closure, canopy height and vegetation heterogeneity (expressed as the Shannon-Wiener diversity of canopy cover at heights interval of 1, 2, 4, 16, 32, and 64 meters) were measured at nine points on each plot and averaged (Ferger *et al*. 2014). The VSI value of each plot was then computed as the mean Euclidian dissimilarity of each plot’s vegetation structure relative to the vegetation structure of plots with undisturbed vegetation at that elevation (Peters *et al.,* 2019). The four environmental variables (MAT, MAP, BRI and VSI) were only weakly correlated across the 44 plots (r < 0.5 in all cases), except for a moderate positive relationship between MAT and BRI (n = 44 plots, r = 0.64, p<0.05).

### 2.6 TReM diversity

We calculated TReM diversity at the level of individual trees and for each plot. At the tree level, we calculated the frequency of each TReM type on every surveyed tree by summing the occurrences of TReMs across each of the one-meter segments of the vertical transect. This resulted in a matrix containing the frequency of all 44 TReM types on every sampled tree.

Based on this frequency matrix, we computed the Shannon diversity index controlling for sampling effort. To this end, we used the number of segments of the vertical transects as a measure of sampling effort (Fig. 1b) and calculated a rarefied estimate of Shannon diversity of microhabitats for every single tree (R package iNEXT, function “estimateD”, sampling coverage = 0.85) (Hsieh *et al*. 2016). At the plot level, we summed TReM occurrences for all the surveyed trees within that plot. Based on these data, we computed a rarefied Shannon diversity using the total number of transect segments as a measure of sampling effort for each plot. Rarefaction was performed so that we would not systematically estimate a higher diversity of habitats in plots with taller trees or a lower diversity on plots on which less than five trees could be sampled, as we always surveyed a maximum of five trees per plot.

### 2.7 Statistical analysis

We tested the hypothesis that larger and architecturally more complex trees hosted a higher TReM diversity by fitting a linear mixed model that related tree-level, rarefied Shannon diversity to each tree’s DBH, height of the first branch, and to species-level wood density. We included plot identity and tree species identity as random factors, because multiple trees were sampled on each plot. While we surveyed a total of 180 individual trees on the field, this analysis uses data from 148 trees, as not all trait measurements were available for all tree individuals.

To test the hypothesis that climate and human impact affect TReM diversity at the plot level, we fitted a linear model that related the plot-level, rarefied Shannon diversity to MAT, MAP, BRI, and VSI. For this analysis we were able to use data from all 180 surveyed trees. For both models, we performed model selection across all possible combination of predictor variables (main effects only) using the function “dredge” from R package MuMIn (Bartoń 2022) using the second order Akaike’s Information Criterion (AICc) to select the best model. For the model addressing the second hypothesis, it was not possible to choose a single best model as three models had an AICc difference lower than two units. In this case, we performed model averaging across all the models that were within a range of two AICc units relative to the best model.

To test whether the composition of TReM assemblages depended on the environmental conditions on each plot, we employed non-metric multidimensional scaling (NMDS) to quantify changes in TReM composition among plots. We used the R package “vegan” (Oksanen *et al*. 2022) to calculate the Bray-Curtis dissimilarity between all pairs of plots based on TReM frequencies, which were standardized via a Wisconsin double standardization, and then applied NMDS scaling to two axes (function “metaMDS”). We tested how microhabitat composition was related to MAT, MAP, BRI and VSI by performing an environmental vector fit onto the NMDS ordination (n=10.000 permutations, function “envfit”, R package “vegan”) (Oksanen et al. 2022).

## 3 Results

We surveyed a total of 180 trees belonging to 41 plant species distributed over 29 plant families (see the plant species list in **Table S1**). We encountered 43 out of the 44 TReM types from our catalogue. The most abundant TreMs were moss patches (1580 occurrences), lichen patches (1026 occurrences) and bark microsoil (759 occurrences). The rarest TreMs were the witch’s broom (1 occurrence), semi-open trunk rot-holes (2 occurrences) and bird foraging excavations (3 occurrences).

### 3.1 TReM diversity at the tree level

At the individual tree level, diameter at breast height and height of the first branch were the most important predictors of TReM diversity, whereas wood density was not significantly associated with TReM diversity (Table 1a). TReM diversity increased strongly with DBH, showing a saturating trend with increasing DBH (Fig. 2a). Our model predicted that trees with a DBH of 40 cm on average harboured an effective diversity of six TReMs, while trees with a DBH of 80 cm harboured more than seven TReMs (Fig. 2a). Conversely, the height of the first branch was negatively correlated with TReM diversity (Fig. 2b). Our model shows that trees with a first branch at a height of two meters hosted on average eight TReMs, while trees branching at 20 m hosted about five TReMs.

**Figure 3.**
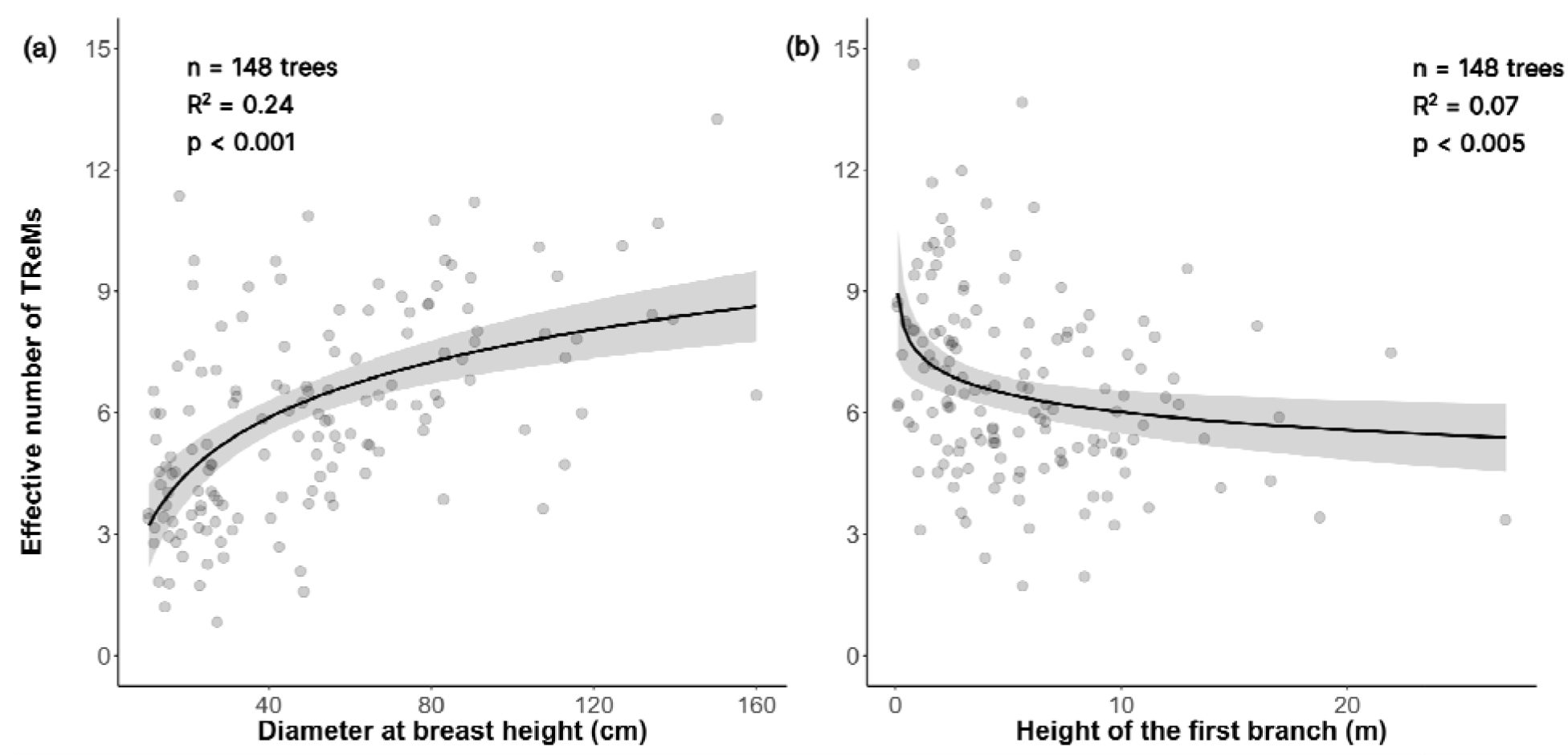
Partial residual plot showing the associations of TReM diversity with DBH and height of the first branch. The Y axis shows the effective number of TReMs in relationship to (a) the diameter at breast height and (b) the height of the first branch. The grey shading represents the 95% confidence interval, while the dots represent the individual trees (n=148). Partial residual plots show the association between response and predictor variable controlling for the effect of other predictors included in the statistical model (see Table 1a for all model coefficients). Shown relationships are non-linear because the values shown are the exponential of the Shannon diversity, which corresponds to the effective number of TReM types on each individual tree (Jost 2006).

### 3.2 TReM diversity at the plot level

MAT and BRI were the most important environmental variables driving TReM diversity at the plot level. TReM diversity increased with increasing temperature (Fig. 3a). BRI was negatively associated with TReM diversity so that human impacts decreased TReM diversity at the plot level (Fig. 3b). VSI and MAP did not show significant relationships with TReM diversity.

**Figure 4.**
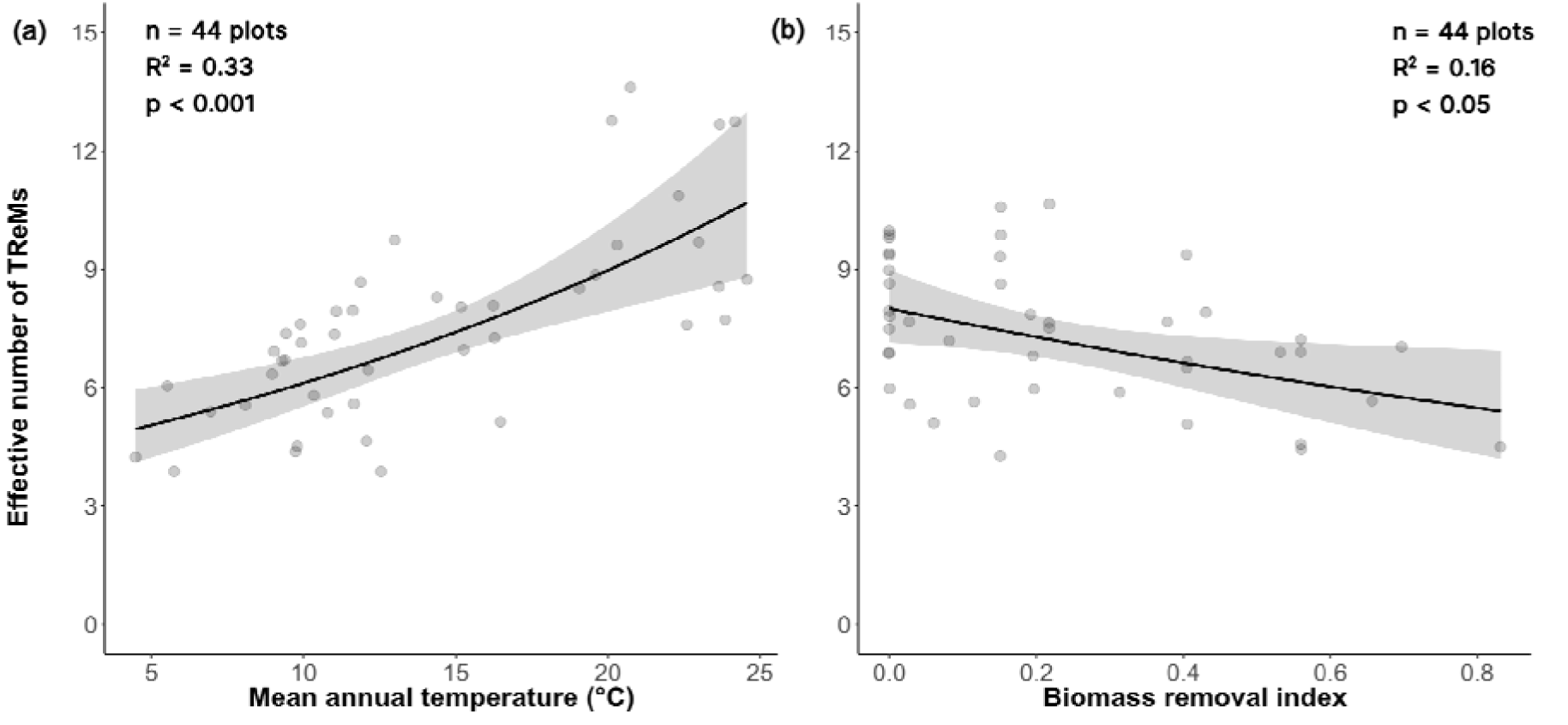
Partial residual plots of the effect of environment on TReM diversity. The Y axis shows the effective number of TReMs in relationship to (a) MAT and (b) BRI measured at the plot level. The light grey shading represents the 95% confidence interval and the dots represent plots. Partial residual plots show the association between response and predictor variable controlling for the effect of other predictors included in the statistical model (see Table 1a for all model coefficients). Shown relationships are non-linear because the values shown are the exponential of the Shannon diversity, which corresponds to the effective number of TReM types on each plot (Jost 2006).

**Table 1.**
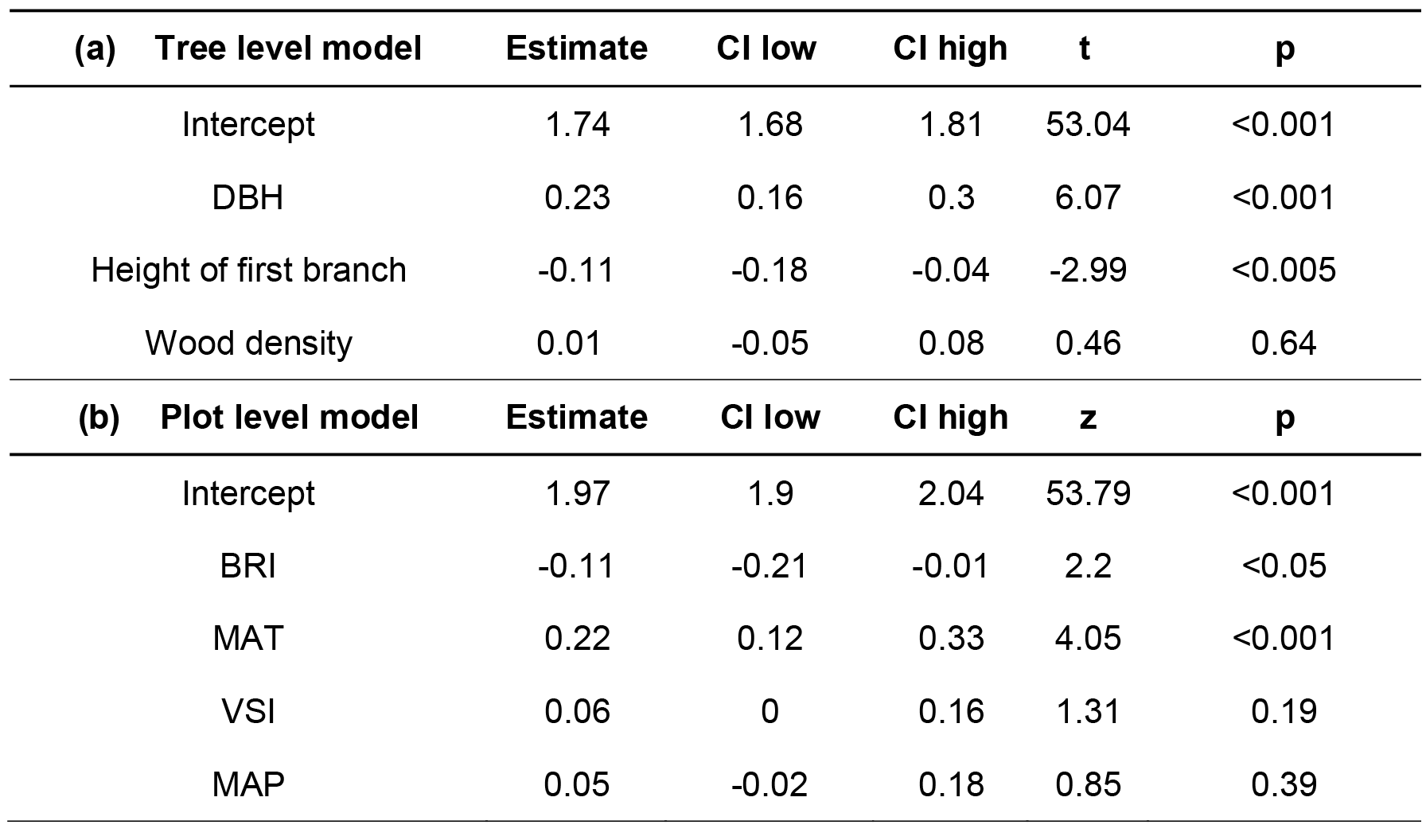
Associations between the diversity of TReM types (a) at tree level and (b) plot level. Given are the results of the linear models relating TReM Shannon diversity to variables describing (a) tree-level and (b) plot-level variability. Parameter estimates are shown in the second column, followed by lower and upper confidence intervals in the third and fourth columns. Test statistics and p-values are given in the last two columns. The tree-level model included random-intercept effects of plot and tree species identity. The plot-level model is based on model averaging across the three best models.

| (a) Tree level model | Estimate | CI low | CI high | t | p |
| --- | --- | --- | --- | --- | --- |
| Intercept | 1.74 | 1.68 | 1.81 | 53.04 | <0.001 |
| DBH | 0.23 | 0.16 | 0.3 | 6.07 | <0.001 |
| Height of first branch | -0.11 | -0.18 | -0.04 | -2.99 | <0.005 |
| Wood density | 0.01 | -0.05 | 0.08 | 0.46 | 0.64 |
| (b) Plot level model | Estimate | CI low | CI high | z | p |
| Intercept | 1.97 | 1.9 | 2.04 | 53.79 | <0.001 |
| BRI | -0.11 | -0.21 | -0.01 | 2.2 | <0.05 |
| MAT | 0.22 | 0.12 | 0.33 | 4.05 | <0.001 |
| VSI | 0.06 | 0 | 0.16 | 1.31 | 0.19 |
| MAP | 0.05 | -0.02 | 0.18 | 0.85 | 0.39 |

### 3.3 TReM composition

TReM composition changed along climatic gradients and in response to human impact (Fig. 4). TReM composition was generally more similar within than among ecosystem types (R^2^ = 0.69, p<0.001) and within forested than non-forested ecosystems (R^2^ = 0.36, p<0.001) (Fig. 4). This turnover in TReM composition was associated with differences in rainfall (R^2^ = 0.69, p<0.001) and temperature (R^2^ = 0.49, p<0.001), while biomass removal was significantly, but more weakly associated with microhabitat composition (R^2^ = 0.2, p<0.01).

**Figure 5.**
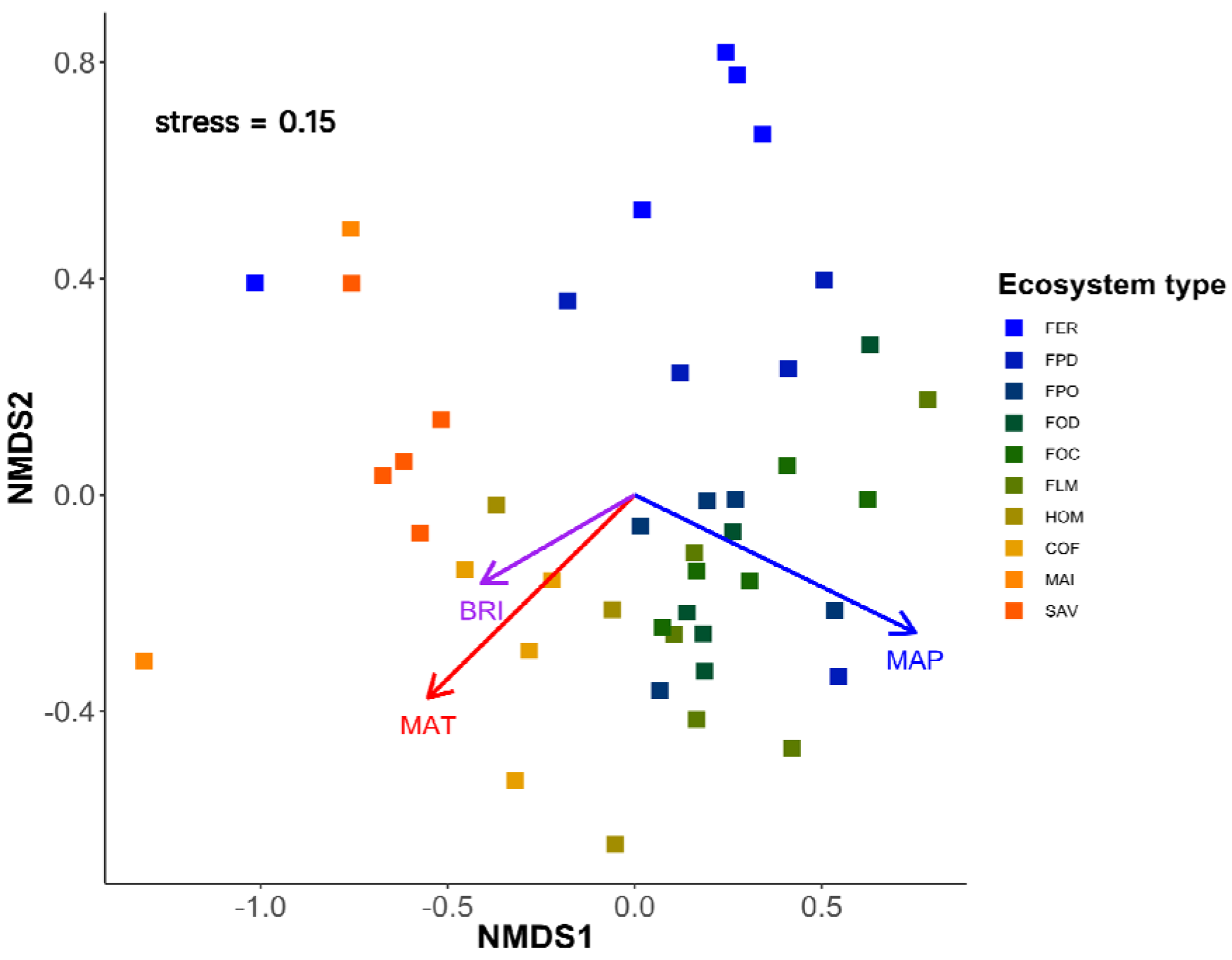
Differences in TReM composition across ecosystem types. Each coloured square represents a plot belonging to one of the ten ecosystem types in which trees were surveyed. Ecosystem types are indicated by the following abbreviations: FER, *Erica* forest; FPD, disturbed *Podocarpus* forest; FPO, *Podocarpus* forest; FOD, disturbed *Ocotea* forest; FOC, *Ocotea* forest; FLM, lower montane forest; HOM, traditional homegarden; COF, coffee plantation; MAI, Maize plantation; SAV, savanna. Closer squares are more similar in TReM composition than squares farther apart. Arrows show the direction of change of the environmental variables that are significantly related to TReM turnover.

## 4. Discussion

This study is the first comprehensive assessment of TReM diversity along a large environmental gradient in the tropics. We found that large trees were particularly important for fostering TReM diversity. Along the studied environmental gradient, MAT had a positive effect on TReM diversity, while human impact had a negative impact. TReM composition was systematically related to both climatic differences and human impact. This work shows that TReMs can be used as sensitive indicators of microhabitat diversity and composition both at the level of individual trees and across large environmental gradients.

### 4.1 TReM diversity at tree level

At the tree level, the strongest predictor of microhabitat diversity was diameter at breast height. This finding is in accordance with patterns observed in temperate forests (Asbeck *et al*. 2021). Diameter at breast height is one of the most widely used measurements of tree size in forestry, because it is easy to record in a standardised way (Blanchard *et al*. 2016). Trees with higher DBH are generally taller and have a larger crown area, and these allometric relationships have been confirmed across multiple tropical forests (Blanchard *et al.,* 2016). This indicates that taller trees with a broader canopy constitute a larger habitat patch for the formation of TReMs. In addition, trees with larger DBH are generally older, which means that they are more likely to show structural changes and damages, such as limb and stem breakages, cracks and bark loss. Moreover, older trees likely host standing deadwood such as dead branches, or rotten areas where cavities are more likely to form naturally (Larrieu *et al*. 2018). This has important implications in terms of conservation and forestry management, as it underlines the importance of large veteran trees as hotspots of biodiversity (Kozák *et al*. 2023; Lindenmayer 2017). Our findings demonstrate that DBH is a cost-effective indicator to measure TReM diversity in forests. In this study, we did not assess TReM diversity on snags, because climbing such trees safely is challenging and requires additional safety measures (Anderson *et al*. 2015). Snags are, however, known to be important in providing TReMs (Paillet *et al*. 2017), and future studies should attempt to include information on how they contribute to microhabitat diversity in tropical forests.

In addition to tree size, we found that the height of the first branch was negatively associated with TReM diversity, indicating that trees with lower first branches tend to have more TReMs. The height of the first branch is generally considered an indicator of a tree’s growth strategy. Trees that branch out at greater heights are likely large leaved trees which show a low degree of ramification (King 1998). These species can be expected to have a less complex canopy architecture which support a lower number of TReMs. While the tree growth form has not been the focus of previous TReM studies, mostly because the diversity of growth forms is lower in temperate than tropical forests, it might be interesting to study such relationships in more detail in tropical ecosystems that are characterised by a high number of tree species and a wide variety of growth strategies.

### 4.2 TReM diversity along the environmental gradient

The diversity of TReMs was positively associated with MAT, so that more microhabitat types were found in warmer areas of the mountain. It is important to consider that several types of TReMs are constituted by living organisms, e.g., epiphytes like ferns, orchids, vines, while other TReMs are produced by living organisms, such as in the case of nests or insect galleries. These types of TReMs had lower incidences at higher elevations, likely due to environmental filtering of its associated organisms. On Kilimanjaro, like in other regions, species richness of plants and animals decreases with elevation and decreasing temperatures (Peters *et al*. 2016). This suggests that temperature is both the major driver of biodiversity and TReM diversity on the mountain.

TReM diversity was negatively affected by human impacts. On Kilimanjaro plant biomass is harvested primarily as a source of timber, fuel or fodder. Timber extraction, in particular, has resulted in changes in the size distribution (i.e., lower mean DBH of trees) in the disturbed plots of *Ocotea* and *Podocarpus* forests. In addition, human impact due to agricultural activities at the lower elevations, e.g., in coffee plantations and the traditional Chagga homegardens, led to a reduction in the number and diversity of trees present in cultivated plots (Hemp 2006). Because large veteran trees are particularly important sources of TReMs, the logging of such trees is directly related to the lower TReM diversity in plots that have high levels of biomass removal. This is in accordance with studies carried out in temperate ecosystems, where managed forests have been found to have lower TReM diversity than old-growth forests (Asbeck *et al*. 2019). Studies of TReMs can therefore be an effective tool to quantify how human impacts shape the habitat heterogeneity provided by forest ecosystems.

### 4.3 Changes in TReM composition

The composition of TReM assemblages in plots was dictated primarily by climate and, to a lesser extent, by human impact. The strongest climatic determinant of TReM composition was rainfall, although it was not significantly associated with TReM diversity. According to our analysis, forested plots clustered together and were distinct from non-forested plots in their TReM composition. Foliose lichens, mosses, and bark microsoil were the most recorded TReMs in this survey and were by a large extent observed in forested plots with high rainfall. Foliose lichens and mosses are known to thrive in conditions of elevated rainfall in the tropics (Benzing 1998) and the highest abundance of ferns and other epiphytes occurs in the more rainy areas of Kilimanjaro (Hemp 2001, 2011). In addition, rainfall is known to be positively associated with the presence of TReMs like limb breakage, stem breakages and other kinds of tree structural damages in tropical forests (van der Meer & Bongers 1996). High levels of precipitation can further trigger tree falls that subsequently damage neighbouring trees and subject tree crowns to strong forces due to high water loads. Similarly, high humidity levels promote accelerated rotting of wood and foster cavity formation (Lindenmayer *et al*. 1993) and are likely to contribute to the distinct TReM communities in the montane forests. Conversely, some TReMs were associated to non-forested plots: the fire damage TReM, for example, was recorded solely in savanna plots, as fire events are frequent in this dry ecosystem. Very large cavity types, like chimney trunk rot holes, and disease related TReMs like decayed cankers occurred mostly in non-forested plots like homegardens and coffee plantations. This might occur because trees in these plots are pruned to harvest fodder and might be exposed to pathogen infections (Mollel *et al*. 2017).

### 4.4 TReMs as a suitable indicator system in the tropics

Our findings demonstrate that TreMs are sensitive indicators of environmental gradients driven both by climatic factors and human impact. This suggests that TReMs are a valuable indicator system for tropical ecosystems, in which they might be particularly useful given the high diversity of species in these ecosystems (Barlow *et al*. 2018) and the difficulty to assess this diversity with cost-effective measures (Schmeller *et al*. 2017). We can speculate that TReMs function as useful biodiversity indicators on Kilimanjaro, because comprehensive biodiversity surveys carried out along this same gradient showed that both the diversity of vertebrate and invertebrate taxa responded similarly to the same set of climate and human impact variables (Peters et al., 2016, 2019). This suggests that TReM diversity is driven by similar mechanisms as biodiversity. There is, however, a need for more studies to investigate the specific relationships between TReMs and biodiversity, especially in the tropics.

Our quantitative survey of the abundance of TReMs on about 200 individual trees required a high number of man hours, as ascending a 30-meter tree can require up to four hours. While TReM surveys from the ground can cover a greater number of trees faster, they might not be able to detect TReMs in the higher areas of the tree canopy, especially under complex tropical canopies (Martin *et al*. 2022; Paillet *et al*. 2015). For future studies, we suggest to test the use of small unmanned aerial vehicles (UAVs) with image recording capabilities that could provide similarly detailed surveys, as they have been successfully employed to monitor epiphytes or to sample arthropods from the forest canopy (Krasylenko *et al*. 2023; Madden *et al*. 2022).

## 5. Conclusions

In this paper, we present the first study of TReMs along a broad climate and human impact gradient on a tropical mountain. We have shown that both the diversity and composition of TReMs are sensitive to changes in climate and human impact across the studied ecosystems. These findings demonstrate the applicability of the TReM concept as a highly sensitive monitoring tool to determine how changes in climate and human impact affect microhabitat availability and composition in the tropics. We believe that our approach to quantify TReM diversity can constitute a cost-effective monitoring tool to be employed in highly diverse ecosystems at tropical latitudes.

## 6. Acknowledgments

This work was funded by the Deutsche Forschungsgemeinschaft (DFG) within the research unit Kili-SES (FOR 5064; grant numbers: SCHL 1934/4-1, HE2719/14-1. We thank the COSTECH, TAWIRI and TANAPA authorities for providing the permits to conduct fieldwork on Kilimanjaro (permit no: 2022-307-NA-2021-094). We thank the staff at the research station of Nkweseko (Moshi, Tanzania) for hosting us and supporting our fieldwork. Special thanks go to Raymond Vitus, Frederick Issaack and Esrom Nkya who assisted with the TReM survey.

