## supplementary for "Changes in the diversity and composition of tree-related microhabitats across climate and human impact gradients on a tropical mountain"

**Supplementary Information**

**1. TReM typology for this study**

We detail below the typology used for identifying TReMs in this study, which is based on the typology developed by (Larrieu *et al.* 2018) but has been subjected to some changes to be suitable for tropical ecosystems and tree-climbing approach (Nußer et al., 2024). The typology used here consists of 6 TReM forms and 44 TReM types.

| **TReMs by grouped by form** | **Frequency** |
| --- | --- |
| **Cavities** | |
| Woodpecker, barbet, tinkerbird nesting cavities | 8 |
| Trunk base rot hole | 15 |
| Trunk rot hole | 96 |
| Semi-open trunk rot hole | 2 |
| Chimney trunk base rot hole | 4 |
| Chimney trunk rot hole | 15 |
| Hollow branch | 50 |
| Insect galleries and bore holes | 166 |
| Dendrotelm | 14 |
| Bird foraging excavations | 3 |
| Bark lined concavity | 53 |
| Root buttress concavity | 53 |
| **Injuries** | |
| Bark loss | 149 |
| Fire scar | 38 |
| Bark shelter | 289 |
| Bark pocket | 249 |
| Stem Breakage | 18 |
| Limb breakage | 198 |
| Crack | 97 |
| Lightning scar | 0 |
| Fork split at insertion | 3 |
| **Exudates** | |
| Sap run | 8 |
| Heavy resinosis | 7 |
| **Deadwood** | |
| Dead branches | 539 |
| Dead top | 8 |
| Remaining broken limb | 109 |
| **Diseases** | |
| Witch’s broom | 1 |
| Epicormic shoots | 9 |
| Burr | 67 |
| Decayed canker | 24 |
| **Epiphytes and epiphytic structures** | |
| Polypores | 17 |
| Other fungi | 3 |
| Ferns | 531 |
| Orchids | 16 |
| Lianas & Vines | 214 |
| Mistletoes | 24 |
| Bryophytes | 1580 |
| Foliose lichens | 1026 |
| Vertebrate nests | 6 |
| Invertebrate nests | 18 |
| Bark microsoil | 759 |
| Crown microsoil | 82 |
| Other epiphytes | 447 |
| Dead leaves frill | 9 |

**Table S1 – Tree species list**

| **Species** | **Family** |
| --- | --- |
| *Acacia sp.* | Fabaceae |
| *Agarista salicifolia* | Ericaceae |
| *Agauria salicifolia* | Ericaceae |
| *Albizia chinensis* | Fabaceae |
| *Albizia gummifera* | Fabaceae |
| *Albizia schimperiana* | Fabaceae |
| *Boswellia neglecta* | Burseraceae |
| *Casearia battiscombei* | Flacourtiaceae |
| *Cassia siamea* | Fabaceae |
| *Combretum molle* | Combretaceae |
| *Cordia africana* | Boraginaceae |
| *Cupressus lusitanica* | Cupressaceae |
| *Cyathea manniana* | Cyatheaceae |
| *Dendrosenecio kilimanjari ssp. cottonii* | Asteraceae |
| *Dombeya rotundifolia* | Malvaceae |
| *Entandrophragma excelsum* | Meliaceae |
| *Erica excelsa* | Ericaceae |
| *Erica trimera* | Ericaceae |
| *Ficus sur* | Moraceae |
| *Garcinia tansaniensis* | Clusiaceae |
| *Grevillea robusta* | Proteaceae |
| *Hypericum revolutum ssp keniense* | Hypericaceae |
| *Ilex mitis* | Aquifoliaceae |
| *Lannea schimperi* | Anacardiaceae |
| *Macaranga kilimanjarica* | Euphorbiaceae |
| *Maesopsis eminii* | Rhamnaceae |
| *Moringa oleifera* | Moringaceae |
| *Newtonia buchananii* | Fabaceae |
| *Ocotea usambarensis* | Lauraceae |
| *Olea africana* | Oleaceae |
| *Ozoroa insignis* | Anacardiaceae |
| *Persea americana* | Lauraceae |
| *Pittosporum spec* | Pittosporaceae |
| *Podocarpus latifolius* | Podocarpaceae |
| *Polyscias fulva* | Araliaceae |
| *Prunus africana* | Rosaceae |
| *Schefflera volkensii* | Araliaceae |
| *Sclerocarya birrea* | Anacardiaceae |
| *Syzygium guineense* | Myrtaceae |
| *Tabernaemontana stapfiana* | Apocynaceae |
| *Xymalos monospora* | Monimiaceae |
| *Ziziphus mucronata* | Rhamnaceae |
